# DynaMiCs – Dynamic cell-type deconvolution ensembles for Mapping in mixed Conditions

**DOI:** 10.1101/2025.05.08.652868

**Authors:** Nicole Seifert, Laurenz Engel, Jana Tauschke, Thomas Sterr, Jakob Demmer, Malte Mensching-Buhr, Dennis Völkl, Sushma Nagaraja Grellscheid, Tim Beissbarth, Franziska Görtler, Helena U. Zacharias, Michael Altenbuchinger

## Abstract

Single-cell techniques facilitate the molecular analysis of individual cells, providing insights into cellular diversity, function, and the complexity of biological systems. However, their application is typically limited to small-scale studies involving individual or a few dozen samples, as a consequence of costs and experimental requirement. This complicates the inference of robust conclusions about populations. Bulk transcriptomics offers cost-efficient measurements with low experimental requirements. However, the cellular resolution is lost and only a complex linear combination of signals from multiple cells is observed. Thus, gene expression changes cannot be attributed to individual cells or cell populations.

Cell-type deconvolution methods infer cellular compositions from bulk transcriptomics data. State-of-the-art approaches use single-cell data to build molecular reference profiles and identify powerful cell-type markers for improved deconvolution. In this context, we propose Dynamic cell-type deconvolution ensembles for Mapping in mixed Conditions (DynaMiCs) for the integration of single-cell and bulk transcriptomics data. Specifically, DynaMiCs dynamically extracts information from single-cell experiments to (1) provide more accurate estimates of cellular compositions, and (2) establish a mapping between bulk and single-cell data. Consequently, DynaMiCs enables the investigation of how cell populations change in both quantity and molecular characteristics between different phenotypes, informed by single-cell experiments.

## 1 Introduction

Single-cell RNA sequencing provides a snapshot of individual cells’ transcriptomes, enabling detailed insights into cellular diversity and function. In contrast, traditional bulk techniques lack cellular resolution, as they measure averaged gene expression across complex mixtures of many cell types. This masks cell-specific contributions and prevents direct attribution of gene expression changes to distinct populations. Consequently, bulk studies provide only a limited picture of cellular heterogeneity. However, single-cell techniques remain expensive and experimentally demanding, limiting their use to relatively small cohorts[1]. This limits single-cell experiments to few selected samples, although with increasingly high throughput, capturing 100,000s to millions of individual cell profiles. As a consequence, conclusions about populations, such as diseased patients, remain challenging as those require large-scale studies, involving many biological replicates. Furthermore, experiments can be confounded by batch effects and inherent cellular variability, complicating the integration of different single cell experiments [2]. Therefore, even if costs and experimental requirements are not a limiting factor, it remains challenging to draw conclusions from large-cohort single-cell data.

Cell-type deconvolution approaches emerged as reliable tools for the computational inference of cell compositions from bulk transcriptomics data [3]. In typical bulk analyses, it remains unclear whether observed gene-expression changes are the consequence of differential gene regulation or differences in cellular compositions, and revealing the underlying cellular compositions is crucial for discovering the true biological mechanisms.

Most state-of-the-art methods for cell-type deconvolution are reference-based with cell populations represented by reference profiles [4, 5]. These profiles are combined as a weighted sum to resemble the observed bulk, yielding pseudo-quantitative measures of cellular distributions: weights usually serve as a surrogate which correlate with cell number but do not represent the actual number of cells in a mixture [6]. Noteworthy, cellular proportions can be estimated under the assumption that all relevant cell types are represented in the reference data— i.e., there are no hidden contributions from unmodeled or unknown populations [7].

Single-cell data can be leveraged to construct reference profiles by aggregating cell profiles of specific cell types [8]. Furthermore, it was demonstrated that reference-based approaches yield improved estimates when the references are limited to relevant cell markers [8]. Ideally, this would include genes that help to disentangle cellular contributions while exclude those that do not provide useful information. This also provides an objective for machine learning algorithms to improve cell-type deconvolution. Görtler et al. proposed a machine learning framework to weigh genes according to their relevance for cell-type deconvolution, where artificial bulk mixtures from single-cell data serve for model training [6]: the algorithm up-weights genes which improve cell type deconvolution while it down-weights those which do not. Alternative machine learning approaches directly predict cellular proportions without any need for references [9]. Scaden was shown to provide state-of-the-art performance on the same level as the top reference-based approaches [3]. Reference based methods were further adapted to account for common domain issues. For example, models trained on single-cell data from healthy breast tissue have been shown to transfer reliably to cancerous breast tissues [7], when background contributions and tissue adaptation factors are inferred during the deconvolution procedure.

Unlike bulk techniques, single-cell methods provide a detailed molecular characterization of individual cells. Cell populations can be directly defined and characterized via clustering approaches, yielding a rich resource to study condition-related molecular changes in cell populations. This information remains hidden for cell-type deconvolution, although promising recent works, such as the CIBERSORTx [8], TissueResolver [10], ADTD [7], and BayesPrism [11], facilitate estimates of cell-type specific gene expression. These approaches use single-cell data to identify/weigh genes, build cell-type specific reference profiles, or directly as references for deconvolution. Here, we propose DynaMiCs, an ensemble-based cell-type deconvolution framework that systematically integrates single-cell data into bulk analyses through dynamic reference construction and gene re-weighting. Specifically, DynaMiCs (a) provides a machine learning framework to combine single-cell data to content-specific references, (b) optimizes deconvolution via gene re-weighting, and (c) provides a direct mapping from bulk to single-cell data. The latter provides a rationale to study disease-related alterations in cell populations and their developmental trajectories from bulk transcriptomics data.

## 2 Results

### 2.1 DynaMiCs in a nutshell

#### Reference matrices and gene weights

DynaMiCs follows a two-step procedure. First, artificial bulk mixtures simulated from single-cell data are used to optimize cell-type specific references and gene weights. Here, the ground-truth cell compositions serve as optimization target, where two sets of model parameters are trained to refine the deconvolution procedure (Figure 1A-C). The first set are weights **Θ** in a linear combination of single-cell profiles to form cell-type specific reference profiles (Figure 1A,B). The second set are gene weights ***g*** to up- and down-weight gene-specific contributions of the deconvolution equation (Figure 1B). Both parameter sets are optimized jointly to improve deconvolution performance (Figure 1C).

In the following, we illustrate this procedure for an exemplary simulation study. We retrieved single-cell RNA sequencing data derived from blood of 18 COVID-19 patients with a mild (*n* = 9) or severe (*n* = 9) disease course from [12], and simulated artificial bulks as described in the Methods section 4.4. Using these bulks of known cellular composition, we trained gene and profile weights jointly with DynaMiCs.

**Figure 1:**
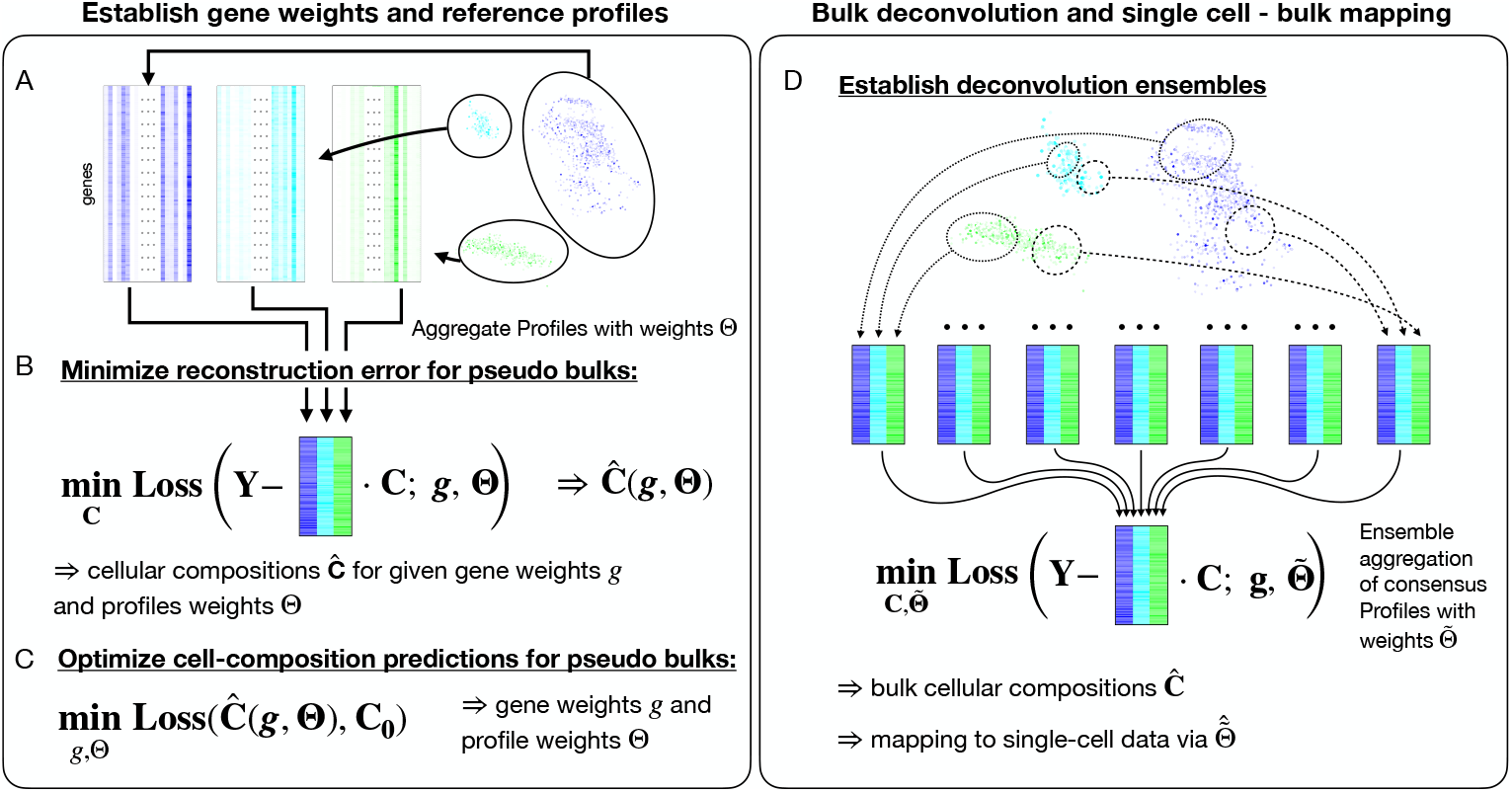
**(A-C)** Illustration of the DynaMiCs algorithm to establish gene weights and reference profiles. **(A)** Single-cell input data with gene expression levels in rows and cells in columns, respectively. Different colors correspond to different cell types. Color intensities illustrate the contribution of individual cell profiles to the consensus profiles in **(B)**. Respective weights are determined via the optimization procedure **(B**,**C)** with **(B)** showing the internal loss quantifying the reconstruction quality of the pseudo bulk profiles in **Y** by a weighted linear combination of reference profiles. The weights **C** = (*c*_*ki*_) can be interpreted as pseudo-quantitative measures of cell distributions, where rows correspond to cell types *k* = 1, …, *q* and columns to different cell mixtures *i* = 1, …, *n*. The reference profiles themselves are defined as a linear combination of single-cell profiles with trainable weights **Θ** = [*θ*^(1)^, …, *θ*^(*q*)^], where *θ*^(*k*)^ contains the weights for cell type *k*. **(C)** Outer loss function quantifying the prediction accuracy, where **C**_0_ contains the ground-truth cellular proportions of the pseudo bulks and **Ĉ** (***g*, Θ**) the estimates derived by optimizing the inner loss in Figure **(B)**. Latter estimates depend on the gene weights ***g*** and the profile weights **Θ**. The outer loss **(C)** thus defines an optimization problem to determine optimal gene and profile weights. **(D)** Ensemble loss to decompose bulk profiles via an ensemble of reference matrices determined by profile weights 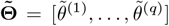, where 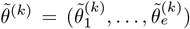 contains the weights of ensemble 1 to *e* of cell type *k*. Thus, ensembles are weighted to resemble the bulks. Since the ensemble profiles themselves are trained via procedure **(A-C)**, this procedure can be used to map bulks to single cell profiles via a combination of the ensemble weights 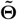 and the single-cell profile weights **Θ**.

Figure 2A shows model performances for each of the captured cell types versus the number of iterations in the optimization procedure, where we measured performance using Pearson’s correlation, comparing the predicted cellular proportions to the ground truth. Some cell types such as B cells, CD4+ and CD8+ T cells can be predicted reasonably well without training (0 iterations) with *r* ∼0.74, *r* ∼ 0.71 and *r* ∼ 0.78, respectively. Nevertheless, performance rises with increasing iterations, converging at approximately *r* ∼ 0.92 for 500 iterations. Predictions for blood cells and gamma-delta (*γδ*) T cells, in contrast, fail without optimization with *r* ∼ 0.19 and *r* ∼0.30, respectively. With increasing iterations, these values rise to *r* ∼0.69 and *r* ∼0.76, respectively. Importantly, blood cells and *γδ* T cells constitute a minor contribution to the bulks with average contributions of 6% and 1%, which might explain the compromised predictions for the naive estimate without optimization. The established profile weights provide a mapping from bulk to single-cell data. DynaMiCs starts with a random initialization of profile weights, as illustrated in UMAP space in Figure 2B for CD4+ and CD8+ T cells (upper and lower row, 0 iterations). These weights become increasingly nuanced for 200 and 500 iterations (second and third column), where the former distributes gene weights across a high number of cells (middle column), while the latter focuses on selective cells (right column).

**Figure 2:**
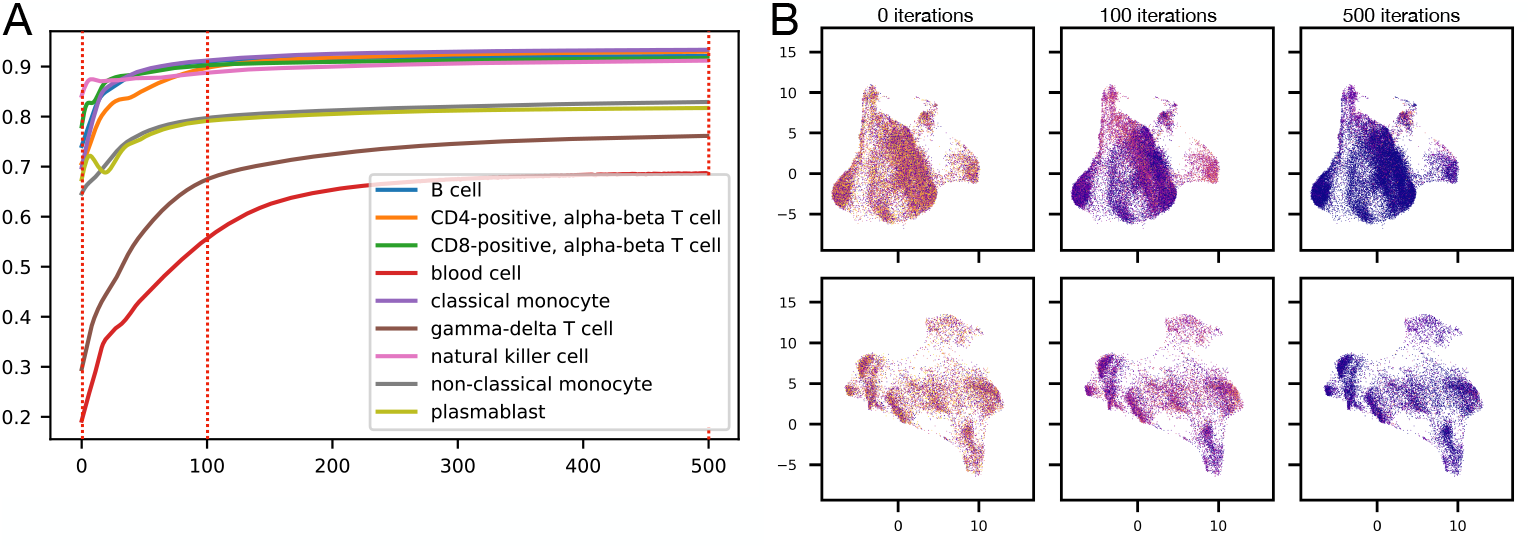
**(A)** Cell-type specific prediction performance versus iterations in the optimization procedure. Performance is measured in terms of Pearson’s correlation, comparing the predicted cellular proportions to the ground truth. **(B)** UMAP plots for CD4+ and CD8+ T cells (upper and lower row). Colors correspond to profile weights from low (blue), average (red), to high weights (yellow).

To assess the biological plausibility of the learned gene weights, we analyzed enrichment of known cell-type markers within the top-ranked genes of reference matrices across training iterations. Genes were ranked by the product of their trained gene weight and normalized expression, and evaluated for enrichment of cell-type markers derived from a publicly available human PBMC reference [13]. Figure 3A–C shows marker enrichment curves for one representative training run (A: without training, B: after 100 iterations, C: after 5,000 iterations), with black ticks marking the ranks of known marker genes. Already after 100 iterations, marker genes are strongly enriched among the top-ranked features, with peak enrichment scores exceeding 0.7. Figure 3D–F visualizes how the evolving gene weights shape the reference matrices during training, with the top 100 genes extracted at 0, 100, and 5,000 iterations, and the position of known marker genes marked with black bars. Before training (Figure 3D), the reference profiles are dominated by highly expressed genes and include only two known markers. Clustering fails to separate cell types meaningfully, for example leading to CD8+ T cells clustering closer to blood cells than to CD4+ T cells, or B cells clustering closer to CD4+ T cells than to plasmablasts. After 100 iterations (Figure 3E), marker frequency increases fivefold compared to the untrained reference (Figure 3D), and biologically meaningful clusters begin to emerge. Notably, while the pan-leukocyte marker B2M drops out of the top 100 genes, its functional equivalent HLA-B — also associated with MHC class I presentation — rises in rank, indicating a refinement rather than loss of biological signal. By iteration 5,000 (Figure 3F), the marker frequency slightly declines but remains markedly higher than the pre-training baseline. Interestingly, more genes with low overall expression begin to populate the reference matrices at later iterations, suggesting that the optimization process increasingly captures subtle, yet informative, transcriptomic features that contribute to cell-type discrimination beyond canonical markers.

The representative training run shown in Figure 3 reflects a consistent trend observed across 100 simulations. Marker frequencies significantly increased within the first 100 iterations and, although they declined by iteration 5,000, remained significantly higher than in the untrained model (Supplementary Figure 5 and Supplementary Table 2). This pattern suggests that DynaMiCs rapidly prioritizes biologically meaningful features early in training, followed by a gradual shift toward incorporating more subtle or complementary signals for more refined cell-type discrimination.

#### Inference – cell-type deconvolution ensembles

Figure 1A-C illustrates the DynaMiCs training procedure to establish gene and profile weights. To infer cellular compositions from bulk data, DynaMiCs uses an ensemble of deconvolution models, each contributing individual gene weights and reference profiles established via steps 1A-C. The basic idea is to provide DynaMiCs with pseudo bulks generated from different subsets of single-cell data (Figure 1D). Then, for each subset, DynaMiCs establishes characteristic cell-type specific consensus profiles. Those are provided as an ensemble of reference profiles, which are subsequently aggregated to resemble the bulks at consideration. This aggregation procedure is built such that the estimated weights **Ĉ** correspond to pseudo-quantitative estimates of cellular proportions. The detailed procedure is described in Methods section 4.2.

**Figure 3:**
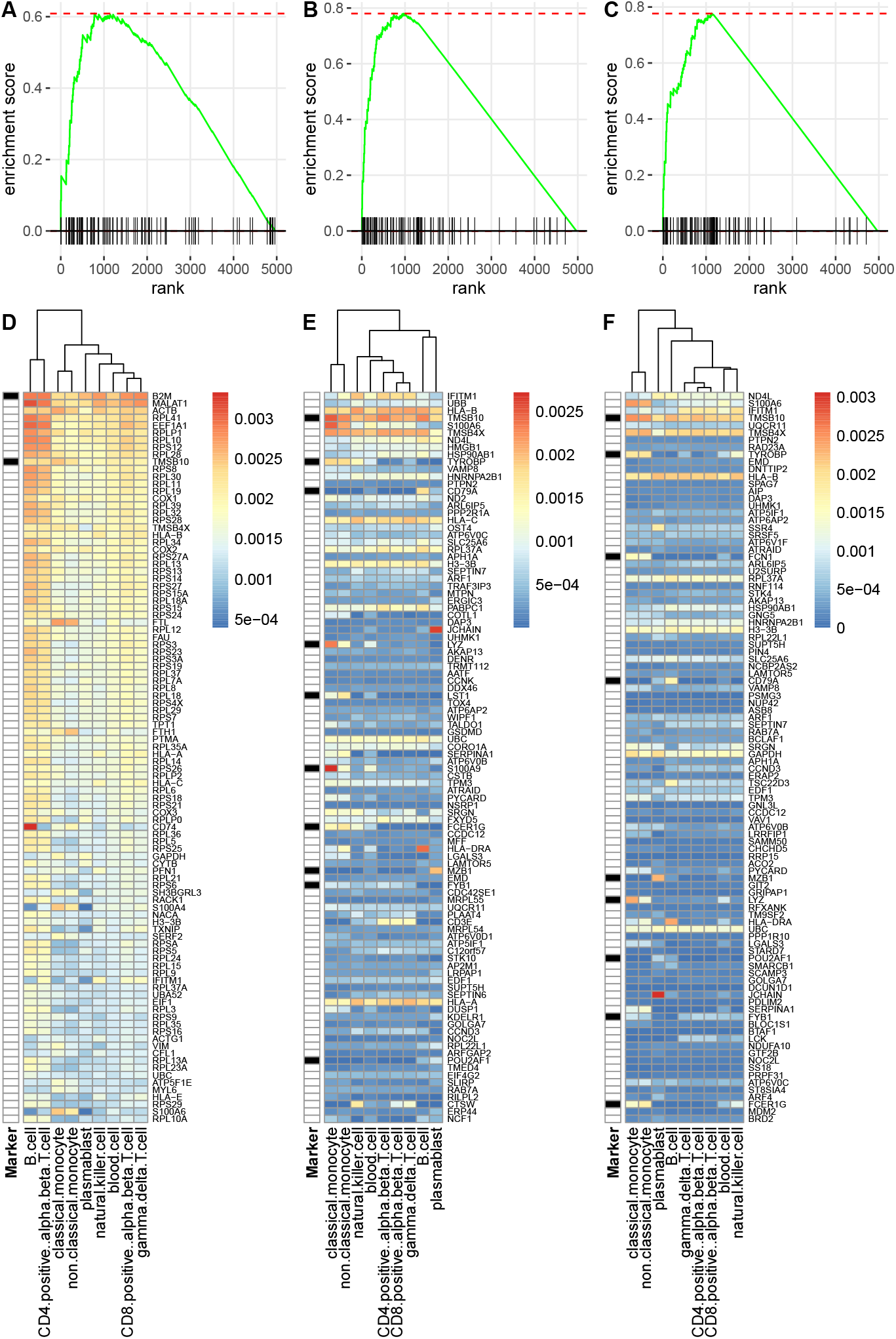
Gene marker enrichment across iterations in the optimization procedure for one representative training run. Genes were ranked by the product of their learned gene weight and normalized expression. Black bars indicate known marker genes and their position in the ranking. **(A-C)** Cumulative distribution of canonical cell-type markers before training (A), after 100 iterations (B), and after 5,000 iterations (C). **(D-F)** Heatmaps of the normalized expression values for the top 100 ranked genes from the reference matrices before training (D), after 100 iterations (E), and after 5,000 iterations (F).

### 2.2 Benchmark analysis

We performed a comprehensive benchmark analysis comparing DynaMiCs against four state-of-the-art deconvolution methods; BayesPrism, MuSiC, Scaden, and SCDC. To build artificial bulks, we downloaded data from the COVID-19 Multi-omics Blood ATlas (COMBAT) Consortium [12], which is a comprehensive resource providing multi-omic data from patients with varying COVID-19 severity. Single-cell RNA-seq data were separated according to patient IDs into a training and test cohort, each containing (*n* = 9) samples of mild and (*n* = 9) samples of critical COVID cases. The training data were then directly used as an input to BayesPrism, MuSiC, and Scaden, as well as for DynaMiCs. For DeconRNASeq, we provided a reference matrix aggregated from the single-cell training data as input. Next, pseudo bulks of known cellular composition were generated from the test cohort by randomly drawing and aggregating 100 single-cell profiles. The latter step was performed such that each bulk mixture contains only single-cell profiles corresponding to one individual patient. This procedure was chosen to ensure that the variability of cellular compositions resembles those of real bulks. The performance in terms of Pearson’s correlation for all competing methods is summarized in Table 1. We observed that DynaMiCs performed best for 6 out of 9 cell types, while BayesPrism, MuSiC, and Scaden each performed best for only one cell type. While some cell types, such as CD8+ T cells, were reasonably well estimated by all methods, performance gains were substantial for less dominant cell populations, such as non-classical monocytes and plasmablasts. For instance, non-classical monocytes were estimated with *r* = 0.69, while the second-best method in this comparison was Scaden, with *r* = 0.42.

**Table 1:** Comparison of cell-type deconvolution approaches. Performance was assessed by calculating the Pearson’s correlation between predicted cellular proportions and the ground truth for each individual cell type. Error intervals correspond to ±1 standard deviation across repeated simulation runs.

| mode | BayesPrism | DynaMiCs | MuSiC | SCDC | Scaden |
| --- | --- | --- | --- | --- | --- |
| B cell | $0.818 \pm 0.031$ | **$0.921 \pm 0.015$** | $0.770 \pm 0.042$ | $0.729 \pm 0.048$ | $0.720 \pm 0.106$ |
| CD4+ T cell | $0.794 \pm 0.055$ | $0.733 \pm 0.068$ | $0.763 \pm 0.051$ | $0.758 \pm 0.053$ | **$0.826 \pm 0.070$** |
| CD8+ T cell | $0.910 \pm 0.025$ | **$0.942 \pm 0.024$** | $0.891 \pm 0.030$ | $0.828 \pm 0.041$ | $0.898 \pm 0.033$ |
| blood cell | $0.275 \pm 0.046$ | $0.251 \pm 0.050$ | **$0.335 \pm 0.039$** | $0.343 \pm 0.026$ | $0.155 \pm 0.108$ |
| cl. monocyte | $0.906 \pm 0.020$ | **$0.949 \pm 0.007$** | $0.890 \pm 0.014$ | $0.903 \pm 0.008$ | $0.898 \pm 0.058$ |
| γ-δ T cell | **$0.701 \pm 0.052$** | $0.451 \pm 0.025$ | $0.606 \pm 0.046$ | $0.653 \pm 0.052$ | $0.431 \pm 0.132$ |
| NK cell | $0.874 \pm 0.025$ | **$0.903 \pm 0.013$** | $0.871 \pm 0.018$ | $0.840 \pm 0.025$ | $0.873 \pm 0.056$ |
| non-cl. mono. | $0.241 \pm 0.066$ | **$0.686 \pm 0.041$** | $0.306 \pm 0.064$ | $0.207 \pm 0.066$ | $0.423 \pm 0.084$ |
| plasmablast | $0.643 \pm 0.047$ | **$0.750 \pm 0.036$** | $0.564 \pm 0.057$ | $0.571 \pm 0.072$ | $0.585 \pm 0.072$ |

### 2.3 Mapping single-cell to bulk transcriptomics with DynaMiCs

DynaMiCs extends conventional cell-type deconvolution algorithms by a mapping between single-cell and bulk transcriptomic data. In a next step, we verified this mapping by repeating the previous simulation study, but now including the COVID-19 phenotype – mild and severe disease course – into the gene and profile weight training procedure. Specifically, we established one deconvolution model (weights and profiles) using the single-cell training data of patients with mild COVID-19 disease course only and one using the data of patients with a critical disease course only. Then, the established DynaMiCs inference algorithm contained two representative reference profiles for each cell type – one for severe and one for critical cases. We then applied this model individually to 100 bulks simulated from the test data and verified whether DynaMiCs correctly selects the reference profile for mild and severe cases for each considered cell type. In other words, we tested whether DynaMiCs can classify COVID-19 mild and severe cases in a cell-type specific way. For this purpose, the corresponding profile weights were translated to classification probabilities by rescaling respective weights to yield an overall sum of one. The corresponding results are shown in Figure 4, where we calculated receiver operating characteristics (ROC) curves and precision recall (PR) curves for each considered cell type. Considering areas under the ROC curves, the best classification was achieved by using profile weights of classical monocytes (area ∼ 0.84), followed by CD8+ and CD4+ T cells (∼ 0.83 and ∼0.82) (Figure 4 left). These results are substantiated by corresponding areas under the PR curve (Figure 4 right). The only cell type which did not contain meaningful information for this classification task were NK cells. Interestingly, the performance was even lower than by chance with an area under the curve of 0.36. One should note, however, that although in total *n* = 100 pseudo bulks were classified for this analysis, the corresponding test data correspond to only *n* = 18 patients, which might explain this erroneous finding. Finally, we verified that this DynaMiCs model also provides correct estimates for cellular compositions. For this purpose, we applied it to pseudo bulks generated by aggregating single-cell profiles of the individual patients from the test data (*n* = 18) with results given in Figure 6. These results are in line with our benchmark study, indicating state-of-the-art performance.

**Figure 4:**
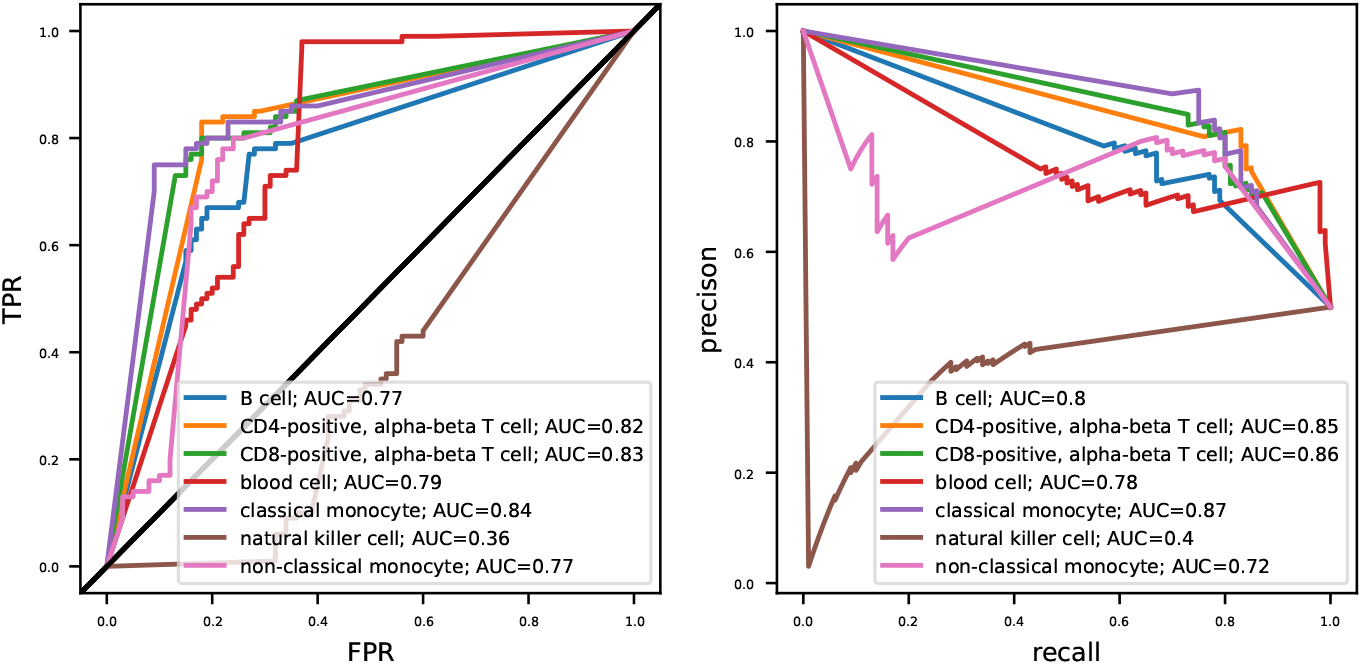
DynaMiCs for cell-type specific bulk classification. Receiver operating characteristics (ROC) curves and precision recall (PR) curves are shown for each considered cell type. Corresponding cell type-specific classification probabilities were extracted from the profile weights established during the DynaMiCs inference step.

## 3 Discussion

Cell-type deconvolution has emerged as an established approach to estimate cellular compositions from bulk transcriptomics data. It has been shown that optimized gene selection and reference profiles can substantially improve deconvolution [8, 6]. For that purpose both prior knowledge and single-cell data can be utilized. Moreover, machine learning approaches play an increasingly pivotal role for cell-type deconvolution, either as direct learning algorithms [9] or to establish cell-type references [6]. In this context, single-cell data are typically used to generate appropriate training data (via pseudo bulks). Recent research has further shown that machine learning can mitigate issues related to domain shifts [14, 7]. Here, the intention is to establish models which are insensitive to the context, meaning that they are trained once and can be reused many times. This, however, might require the adherence to strict SOPs [14]. In this work, we considered a slightly different perspective. A typical workflow is that a restricted set of single-cell data are available to study a specific question, together with a usually sizable set of bulk measurements. The latter are generated to facilitate conclusions on the level of a population, while the single-cell data usually correspond to few patient specimens only (∼ 10 or less). Therefore, we suggest a cell-type deconvolution approach called DynaMiCs, which dynamically integrates single-cell data into the deconvolution procedure. We could demonstrate that DynaMiCs provides state-of-the-art performance, while providing a mapping to single-cell data for downstream analysis. However, there are still limitations. First, it was shown that domain adaptation might require additional algorithmic ingredients. Those comprise potential background contributions and domain adaptation factors which multiply the reference matrix [7]. Moreover, a further future direction could by an extension to spatial omics, where comparable questions are likely of relevance. Still, DynaMiCs extends the scope of cell-type deconvolution algorithms by an integrative mapping between single cell and bulk data. As such, it allows to study how the molecular phenotype of specific-cell populations change between tissues, disease categories and states, from bulk transcriptomics data.

## 4 Methods

### 4.1 Nomenclature

We denote the *j*th column vector of a matrix *A* = (*a*_*ij*_) as 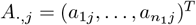 and *k*th row vector of *A* as 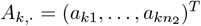. Constraints *a*_*ij*_ ≥ 0 for all *i, j* will be abbreviated as *A* ≥ 0. Estimates of *A* will be indicated by a hat, Â.

### 4.2 Model training – gene weights and reference profiles

#### Single-cell data for model training

DynaMiCs uses labelled single-cell RNA sequencing data for optimized cell-type deconvolution. The basic idea is to generate artificial transcriptomics (ST) data of known cellular compositions from the single-cell data (see subsection “Data simulation”) to train a machine learning algorithm to perform cell-type deconvolution. Formally, our machine learning objective is to provide accurate estimates of cellular compositions, where the trainable model parameters are (1) gene weights and (2) single-cell profile weights, as outlined in the following.

**Gene weights and reference profiles** We first introduce an inner loss function whose optimization returns cellular compositions for a given bulk. Let 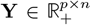 be a matrix containing gene expression profiles of *n* bulks in its columns, where each profile contains gene expression levels for *p* genes. Further, let 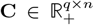 be a corresponding matrix with cellular proportions for *q* cell types in its rows, and 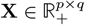 a reference matrix with *q* cell-type specific reference profiles in its columns. Typical cell-type deconvolution approaches use an objective function which quantifies the difference between reconstructed bulks 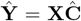 and the observed bulks **Y**. A possible inference strategy could be a non-negative least squares estimate, given by 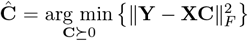. In order to provide more reliable estimates of **C**, we generalize the latter to

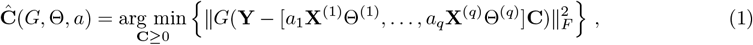

where the 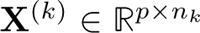 are matrices containing *n*_*k*_ single-cell profiles of cell type *k* in its columns. We further introduce three parameter sets:

- First, *G* = diag(*g*_1_, …, *g*_*p*_) is a matrix with gene weights *g*_*j*_ on its diagonal. Terms corresponding to gene *j* in the sum of squared residuals of (1) are weighted by a factor 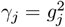. Thus, genes which improve cell-type deconvolution can be up-weighted while those which do not, are down-weighted, respectively.
- Second, *a* = (*a*_1_, …, *a*_*q*_)^*T*^ is a vector of non-negative scalar weights to re-scale the *q* reference profiles. These weights can be adjusted such that the rows of **C** resemble the ground-truth cellular proportions. One should note that row **C**_*j*,·_ multiplies with *a*_*j*_, which contains the cellular proportions for cell-type *j* in the *n* spots. Thus, **C**_*j*,·_ can be scaled by adjusting *a*_*j*_.
- Third, 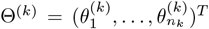 are vectors of non-negative scalar weights 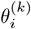, which weigh the contribution of single-cell profile *i* of cell type *k* to the reference profile representing cell type *q*. The vectors Θ^(*k*)^ are collected in Θ = [Θ^(1)^, …, Θ^(*q*)^] and enable the linear combination of single-cell profiles to prototypic references via **X**_·,*k*_ = *a*_*k*_**X**^(*k*)^Θ^(*k*)^.

The estimated cellular proportions Ĉ now depend on the three parameter sets *G*, Θ, and *a*. In order to identify appropriate parameters to facilitate the optimized inference of Ĉ, we state an outer loss quantifying the prediction accuracy,

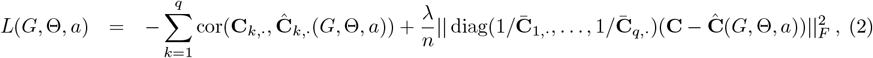

where cor(*x, y*) is the Pearson’s correlation between the vectors *x* and *y* and 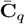 the mean weight of cell type *q*. The DynaMiCs gene weighing and reference profile learning then amounts to the optimization problem

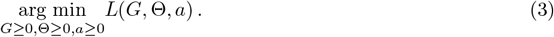

Minimizing − cor(**C**_*k*,·_, *Ĉ* _*k*,·_(*G*, Θ, *a*)), maximizes the Pearson’s correlation between ground-truth cellular proportions **C**_*k*,·_ and respective estimates Ĉ_*k*,·_. In other words, minimizing the first term of *L* maximizes the correlation in predicting cell-type proportions averaged across cell types. Minimizing the second term controls the normalized squared residuals between ground-truth and estimated cellular proportions. This term is included to ensure that the cellular proportions are on the scale of the training data. This is not guaranteed by the first term, as Pearson’s correlation is invariant under the scale of the input vectors, cor(*αx, y*) = cor(*x, y*) for some scalar *α >* 0. The hyper-parameter *λ* calibrates between both optimization objectives and is treated as nuisance parameter. Throughout the analyses presented in this article, we have set *λ* = 0.01.

### 4.3 Inference – cell-type deconvolution

#### Objective function

The former learning procedure infers gene weights and reference profiles for optimized cell-type deconvolution. It also provides an estimate for cellular proportions via optimization problem (1). The full DynaMiCs inference problem is given by

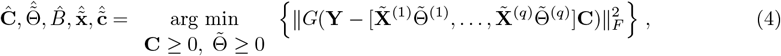

where *G* is determined by the training procedure above. Further, the matrices 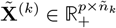 contain *n*_*k*_ proposal reference profiles for cell type *k*, which can be combined to a consensus profile by multiplication with respective weight vectors 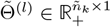. These proposal references themselves can be determined via the DynaMiCs training procedure. Thus, we set 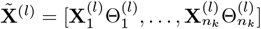, where the subscript gives the training runs 1, …, *n*_*k*_. In other words, we provide a set of pre-aggregated references trained under diverse conditions (e.g., different patients, different diseases) (Fig. 2D), by providing different single-cell training data as input to the DynaMiCs training procedure (Fig. 2A to C).

### 4.4 Real-world data, data simulations and performance evaluations

#### COVID-19 data

We downloaded scRNA-seq data from [12] containing single cell profiles of mild and severe disease courses. Single-cell profiles from one patient were removed (“S00030-Ja003E-PBCa”) as a consequence of a too low cell count number. We constrained the gene space to the 5,000 most variable genes. Cell types with less than 1,000 counts were excluded from the training and labeled as “others” in the test set.

#### Artificial bulks for DynaMiCs

DynaMiCs uses labeled scRNA-Seq data as input to generate artificial bulk mixtures of known cellular composition. These pseudo bulks are used to establish gene and profile weights. For the presented analysis, we established training bulks as follows. For each pseudo bulk, we randomly selected 5 patients and sampled 200 corresponding single-cell profiles. Those were aggregated to yield the pseudo bulk. In total, 10, 000 pseudo bulks were generated for each training run (corresponding to an individual ensemble). For the presented analysis, we used a set of 5 ensembles, meaning that we repeated the procedure of generating pseudo bulks and establishing gene and profile weights five times. In fact, we observed that the performance remains largely unaffected by the specific nature of this training procedure (compare results section), given that a sufficient number of training mixtures were provided for model development.

#### Performance evaluations

For performance evaluations, we separated the COVID-19 data into a training and test cohort. For this, we randomly selected *n* = 9 patients with mild disease and *n* = 9 with critical disease as training data. The remaining *n* = 9 mild and *n* = 9 critical cases were used as test data.

The training data were used to establish deconvolution models, where the data was directly provided to DynaMiCs, BayesPrism, MuSiC, and Scaden, while DeconRNASeq requires a prespecified reference matrix. The latter was generated by aggregating the single-cell training data in a cell-type specific way. DynaMiCs internally performs the model training via pseudo bulks, as outlined above.

Test mixtures were generated patient-wise as follows. We randomly selected single-cell data corresponding to an individual patient and randomly drew 100 single-cell profiles. Those were subsequently aggregated to yield a pseudo bulk of known cellular composition. In total, 1,000 pseudo bulks were generated following this procedure for each test run. In total, 10 test runs were performed for model evaluations.

To assess the robustness of the training and testing procedure, we repeated the same benchmarking setup using varying parameter combinations. Specifically, we varied the number of training runs, the number of training batches and patients per batch, as well as the number of test batches and patients per batch. Supplementary Figure 7 summarizes the resulting performance distributions, indicating that the choice of these parameters has only a minor influence on model performance.

## 5 Acknowledgment

This work was supported by the German Federal Ministry of Education and Research (BMBF) within the e:Med research and funding concept [grant numbers: 01ZX1912A and 01ZX1912C] as well as [grant numbers: 01KD2415A and 01EQ2407A]. Further the work was supported by the German Research Foundation within the TRR247 and the CRU 5002 (426671079).

## 6 Appendix

**Figure 5:**
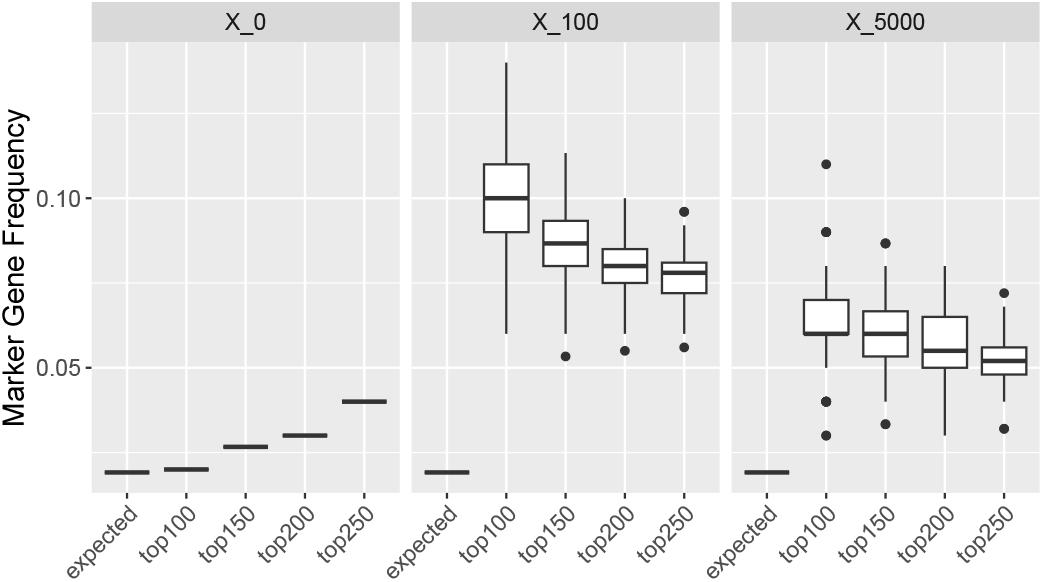
Marker gene frequencies in the top-ranked genes of reference matrices at baseline (X_0), after 100 training iterations (X_100), and after 5,000 iterations (X_5000), compared to the expected frequency in the training data (‘expected’). Genes are ranked by descending score that combines gene weight and normalized gene expression. Boxplots show the distribution across 100 training runs per condition.

**Figure 6:**
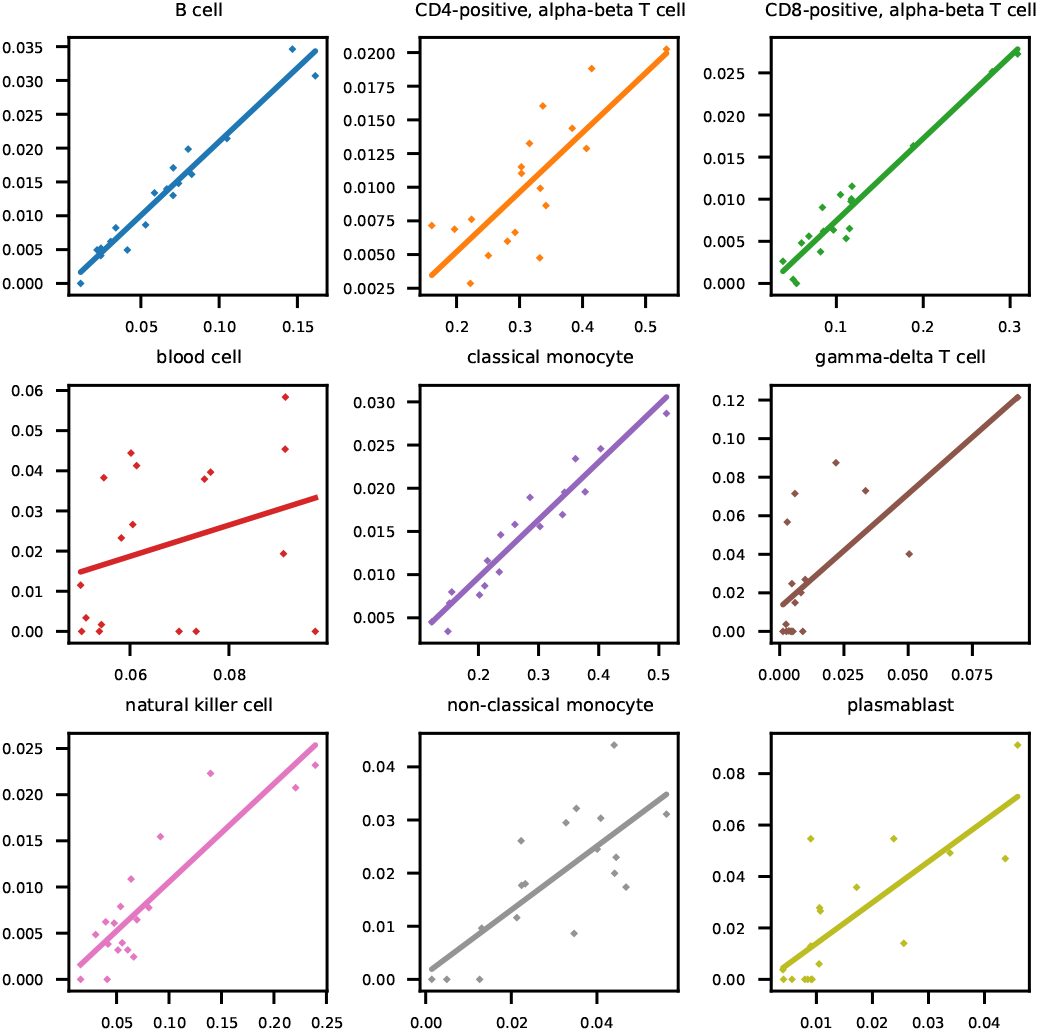
Comparison of DynaMiCs predictions (*y* -axis) with ground-truth values (*y* -axis) for pseudo bulks generated from the single-cell test data (*n* = 18). The colored lines are extrapolations from a least squares estimate.

**Figure 7:**
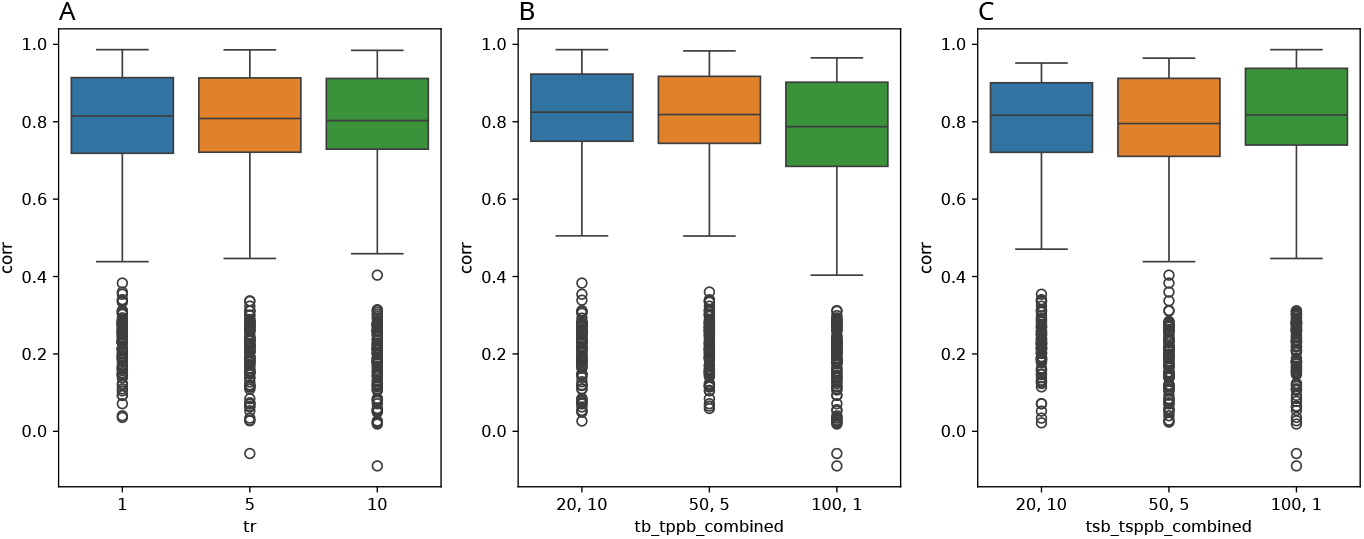
Comparison of DynaMiCs prediction performance under varying training and testing parameters: (A) Number of training runs. (B) Number of training batches and patients per training batch. (C) Number of test batches and patients per test batch. Boxplots show the distribution of Pearson correlation coefficients between predicted and true cell-type proportions.

**Table 2:**
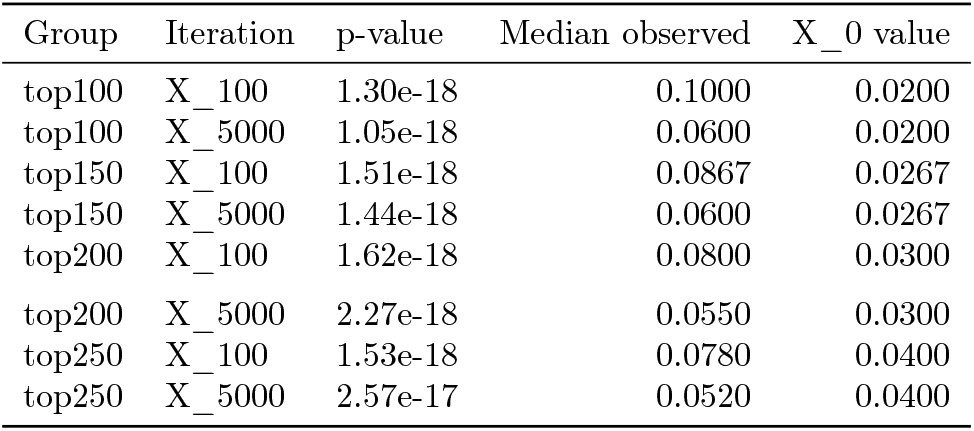
One-sample Wilcoxon test comparing marker gene frequencies in reference matrices after 100 (X_100) and 5,000 iterations (X_5000) to the baseline (X_0 value).

| Group | Iteration | p-value | Median observed | X_0 value |
| --- | --- | --- | --- | --- |
| top100 | X_100 | 1.30e-18 | 0.1000 | 0.0200 |
| top100 | X_5000 | 1.05e-18 | 0.0600 | 0.0200 |
| top150 | X_100 | 1.51e-18 | 0.0867 | 0.0267 |
| top150 | X_5000 | 1.44e-18 | 0.0600 | 0.0267 |
| top200 | X_100 | 1.62e-18 | 0.0800 | 0.0300 |
| top200 | X_5000 | 2.27e-18 | 0.0550 | 0.0300 |
| top250 | X_100 | 1.53e-18 | 0.0780 | 0.0400 |
| top250 | X_5000 | 2.57e-17 | 0.0520 | 0.0400 |

**Table 3:** Comparison of cell-type deconvolution approaches.

|  | test | training | test+training |
| --- | --- | --- | --- |
| B cell | 0.947 | 0.971 | 0.965 |
| CD4-positive, alpha-beta T cell | 0.722 | 0.941 | 0.864 |
| CD8-positive, alpha-beta T cell | 0.923 | 0.844 | 0.884 |
| blood cell | 0.295 | -0.097 | 0.182 |
| classical monocyte | 0.938 | 0.949 | 0.943 |
| gamma-delta T cell | 0.783 | 0.865 | 0.798 |
| natural killer cell | 0.843 | 0.957 | 0.884 |
| non-classical monocyte | 0.710 | 0.881 | 0.828 |
| plasmablast | 0.754 | 0.826 | 0.760 |
| mean | 0.768 | 0.793 | 0.790 |

## Notes

### Competing Interest Statement

The authors have declared no competing interest.

### Summary of Updates

Introduction, first part, has been updated. Methods section, first equation, has been corrected.

